# Mathematical model of the multi-amino acid multi-transporter system predicts uptake flux in CHO cells

**DOI:** 10.1101/2021.04.26.441392

**Authors:** Ashley Sreejan, Mugdha Gadgil, Chetan J. Gadgil

## Abstract

Chinese hamster ovary (CHO) cells express several amino acid (AA) transporters including uniporters and exchangers. Each transporter transports multiple AAs, making prediction of the effect of changed medium composition or transporter levels on individual AA transport rate challenging. A general kinetic model and a simplified analytical expression for the uptake rate is presented. A CHO cell-specific AA transport model, to our knowledge the first such network model for any cell type, is constructed. The model is validated by its prediction of reported uptake flux and amino acid inter-dependencies from experiments that were not used in model construction or parameter estimation. The model defines theoretical conditions for synergistic/repressive effect on the uptake rates of other AAs upon external addition of one AA. This model will help formulate testable hypotheses of the effect of process changes on AA initial uptake, and serve as the AA transport component of kinetic models for cellular metabolism.

## Introduction

Recombinant Chinese hamster ovary (CHO) cell cultures are widely used for the industrial production of therapeutic monoclonal antibodies and several other biopharmaceutical drugs, accounting for more than two-thirds of the approximately USD 200 billion per year market(Walsh, 2018). Amino acids (AAs), in addition to their obvious role in protein translation, are a major contributor to cell mass(Hosios et al, 2016). AA supply at optimal levels is essential to achieve high cell density and productivity(Mulukutla et al, 2017; Salazar et al, 2016a). Commercial production of recombinant therapeutic proteins is increasingly carried out in serum-free and chemically defined culture media. For the amino acid component, medium formulation is usually empirically designed while feeding strategies primarily depend on the estimation or measurement of AA uptake to maintain levels and prevent depletion(Horvat et al, 2020; Sellick et al, 2011; Xing et al, 2011). A quantitative understanding of the effect of AA composition on the individual uptake rates will lead to better-informed medium design and feeding strategies, but is challenging due to the complexity of the amino acid transport system.

Amino acid concentrations may influence not only their own uptake, but that of other AAs. For instance, several studies have shown that uptake competition between cystine and glutamate results in the inhibition of uptake of an AA on the addition of the other(Bannai & Kitamura, 1980; Chen & Swanson, 2003). Glutamate inhibits hybridoma growth due to its inhibition of cystine uptake(Broadhurst & Butler, 2000). Some AAs have an effect on the uptake of other AAs due to their ability to act as an exchange substrate. For instance, intracellular asparagine is a major exchange substrate and regulator of the uptake of AAs such as serine, arginine, and histidine(Krall et al, 2016). Hence, estimating uptake from the AA composition, or predicting a change in the uptake of all AAs due to a change in even a single AA concentration, is challenging. We develop a mechanism-based model of the AA transport system in CHO cells, and use it to predict uptake fluxes from the AA concentrations.

There is an unmet need for a model-based process for optimization of the AA composition profile of CHO cell culture media(Traustason et al, 2019). Most existing models employed for medium optimization do not focus on the transfer of AAs into the cells(Salazar et al, 2016b; Traustason et al, 2019) and hence may not capture the transporter-mediated effect on the observed relationships between AAs. Most CHO cell metabolism models also do not focus on the role of individual transporters in the transfer of AAs. The transport process is modeled as a constant uptake process or assumed to have a mechanism similar to diffusion(Hagrot et al, 2017; Robitaille et al, 2015). Such approaches overlook the complexity of the transport network and its influence on the availability of cellular AAs and transporters. A CHO-specific mechanism-based mathematical model incorporating the known effect of all transporters and AAs would be useful in the estimation of the effect of a specific medium component or transporter perturbation on the uptake flux of all AAs. To this end, we extend previous work on modeling single AA transport in non-CHO cells and formulate a general model, and corresponding simulation software, for AA transport. Using transporter expression data, we customize the extended model for AA transport in CHO cells, and compare its predictions to experimental AA uptake data and reports of amino acid inter-dependencies that were not used in model formulation.

Amino acid transport has mostly been studied at the level of one AAT. However, AA transport in mammalian cells is a complex interconnected process that occurs through a network of transporter systems with multiple mechanisms, viz., uniporters, symporters, and antiporters/exchangers, interlinked through common cargo AAs. Uniporters carry AAs in the thermodynamically favored direction of the concentration gradient. Symporters actively transport AAs driven by an electrochemical gradient (usually of Na+, K+, or Ca+). AA exchangers, also called antiporters(Widdows et al, 2015), exchange one AA from one side of the membrane for another from the opposite side. Each of these AATs can transfer multiple AAs with varying efficacy. Conversely, a single AA can be transported by multiple AATs with different affinities and mechanisms(Bröer, 2008), resulting in a complex combinatorial network of interactions between AAs and AATs. As a result, the transport of one AA depends on the concentration and availability of many AATs. In turn, the availability of a particular AAT for a given AA is dependent on the concentrations and affinities of all AAs being transported by that AAT.

Mathematical modeling has been widely used since the past seven decades to understand the complex process of AA transport, summarized in Table 1. In a few cases, transport has been modeled in a manner analogous to multiphase mass transfer, as the product of an overall mass transfer coefficient and the concentration difference across the membrane(Jacquez, 1973; Weiss et al, 1981). Mechanism based models include explicit AAT-AA binding/dissociation kinetics(Alvarado & Mahmood, 1974; Babcock et al, 1979; Curran et al, 1967; Widdas, 1952), competition(Jacquez, 1961; Jacquez, 1964; Panitchob et al, 2015; Panitchob et al, 2016; Pardridge, 1977; Rosenberg & Wilbrandt, 1957; Sengers et al, 2010), and transport by multiple transporters(Taslimifar et al, 2017). The Taslimafar model(Taslimifar et al, 2017) incorporates both inhibition of transport by competing AAs and additive transport through different transporters to simulate the unidirectional uptake flux in a many-AA many-AAT system. However, this model does not explicitly incorporate AAT-AA binding, or consider transport by AA exchangers.

**Table 1:** Existing mathematical models of amino acid transport. U, S, and E denote uniporter, symporters, and exchangers, respectively, while transport via perfusion/diffusion is denoted by P.† Erythrocytic glucose transporter. ‡ The three terms denote the number of free-carrier, binary-complex, and ternary-complex states.

| No. | Model | AAT types | Explicit AAT-AA binding | Transport direction | No. of states of AATs | No. of AAs or AA pairs | Max no. of AATs of each type | Kinetic asymmetry of AAs |
| --- | --- | --- | --- | --- | --- | --- | --- | --- |
| 1 | Lefevre and Lefevre, 1951 (LeFevre & LeFevre, 1952) | U<sup>†</sup> | Yes | Bi | 1F-1B | 1 Sub. | 1 | Yes |
| 2 | Widdas W. F., 1952 (Widdas, 1952) | U<sup>†</sup> | Yes | Bi | 2F-2B | 2 Sub. | 1 | No |
| 3 | Rosenberg and Wilbrandt, 1957 (Rosenberg & Wilbrandt, 1957) | E | Yes | Bi | 2F-2B | 1 Pair | 1 | Yes |
| 4 | Jacquez J. A., 1961,64 (Jacquez, 1961; Jacquez, 1964) | U+E | Yes | Bi | 2F-2B | 2 AAs | 1 | Yes |
| 5 | Curran et al., 1967 (Curran et al, 1967) | S | Yes | Bi | 2F-2B-2T<sup>‡</sup> | 1 AA | 1 | Yes |
| 6 | Jacquez J. A., 1973 (Jacquez, 1973) | S+P | No | Uni. | NA | 1 AA | 1 | Yes |
| 7 | Stein and Lieb, 1973 (Stein & Lieb, 1973) | U | Yes | Bi | NA | 1 AA | 1 | Yes |
| 8 | Alvarado and Mahmood, 1974 (Alvarado & Mahmood, 1974) | S | Yes | Uni. | 1F-1B-1T<sup>‡</sup> | 1 AA | 1 | Yes |
| 9 | Pardridge W. M., 1977 (Pardridge, 1977) | U | No | Uni. | NA | Many | 1 | Yes |
| 10 | Babcock et al.,1979 (Babcock et al, 1979) | S | Yes | Bi | 2F-2B-2T<sup>‡</sup> | 1 AA | 1 | Yes |
| 11 | Weiss et al., 1981 (Weiss et al, 1981) | S+U | No | Uni. | NA | 1 AA | 1 | Yes |
| 12 | Reinhold and Kaplan, 1984 (L Reinhold & Kaplan, 1984) | U+P | No | Uni. | NA | 1 AA | Many | Yes |
| 13 | Sengers et al., 2010 (Sengers et al, 2010) | S+E+P | No | Bi | NA | Many | Many | No |
| 14 | Panitchob et al., 2015 (Panitchob et al, 2015) | U+E | Yes | Bi | 2F-2B | 2 AAs | 1 | No |
| 15 | Panitchob et. al., 2016 (Panitchob et al, 2016) | U+E+S | Yes | Bi | 2F-2B | 2 AAs | 1 | No |
| 16 | Taslimifar et al., 2017 (Taslimifar et al, 2017) | U | No | Uni. | NA | Many | Many | Yes |
| 17 | Gauthier-Coles et al., 2021(Gauthier-Coles et al, 2021) | U+S+E | No | Bi | NA | Many | Many | Yes |
| **18** | **Proposed Model** | **U+E** | **Yes** | **Bi** | **2F-2B** | **Many** | **Many** | **Yes** |

A series of models that include AAT-AA interactions have been developed by the Sengers group(Panitchob et al, 2015; Panitchob et al, 2016; Sengers et al, 2010; Widdows et al, 2015), primarily to quantify the flux of labeled and unlabeled forms of the same AA. Sengers et al. (2010) describe the net transport flux through symporters by a hyperbolic term as a function of the total external AA concentration. A model for membrane transport, including the competitive effects of substrates, carriers, and the difference in carrier types, was proposed by Panitchob et al.(Panitchob et al, 2015). The underlying structure of the model is the same as the carrier-mediated transport model (Figure 1). This model can explain the action of uniporters as well as exchangers using a single framework of carrier transport. However, since the objective is to model the transport of labeled and unlabeled forms, the interaction of each AA with the carrier was assumed to be identical. Recently, the use of Michaelis-Menten kinetics to calculate ‘homeostasis’ intracellular AA concentrations has been reported(Gauthier-Coles et al, 2021). Exchanger flux is modeled as a multiplication of the unidirectional flux of the exchanged AAs. Seven AA and AATs are included. The model does not aim to capture the transient kinetics of AA uptake, and constrains Km values to preclude AA transport in the absence of a concentration gradient. Analytical expressions for the steady state flux or AA concentrations are not derived.

**Figure 1:**
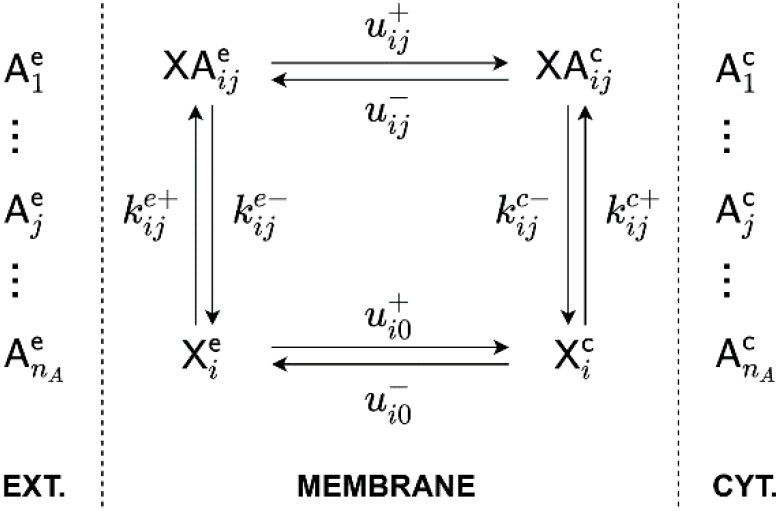
Carrier-mediated transport of n_A_ amino acids. AAs and the carrier are represented by A_j_, and X_*i*_, respectively, and XA_*ij*_ denote the carrier-substrate complex formed by their interaction. Superscripts *e* and *c* represent external and cellular sides of the membrane.

In summary, to our knowledge, none of the mathematical models include the ability to simulate the kinetics of initial uptake of multiple AAs with different transporter types and distinct AAT-AA affinities. Extending the stellar work by the Sengers group, we formulate a computational model and corresponding MATLAB code for AA transport through exchangers and uniporters for multi-carrier multi-substrate systems. For a simplified model, we derive an analytical solution for the steady state AA transport rate for each AA through each transporter. Using data on transporter expression in CHO, we formulate a CHO-specific transport model and predict the uptake flux of AAs in CHO cells under various reported extracellular media conditions. We find that the predictions compare well to experimentally reported fluxes, as well as reported amino acid interdependencies, that were not used in the model formulation. To our knowledge, this is the most comprehensive model for AA transport in CHO cells that is able to quantitatively estimate uptake fluxes for most AAs for various culture conditions and cellular states.

## Results

### A general model and software tool for combinatorial AA transport

We extend the model developed by Panitchob et al. for transport of two AAs to model transport of *n*_*A*_ AAs by one transporter X_*i*_ of *n*_*T*_ transporters by including AA-specific rate constants for binding and bound carrier translocation (Figure 1Figure 1). The carrier (X) can exist in two states X^*e*^ and X^*c*^, corresponding to external and cytosolic surfaces. AAs on one side of the membrane are transported to the other side in the form of a carrier-substrate complex. By disallowing the translocation of free carriers 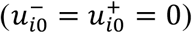, the same scheme can be used for representing an obligatory exchange process.

As described in SI, we have used a single cell as a basis following the methodology of Laufenburger and Linderman(Lauffenburger & Linderman, 1996). AAs levels are expressed as concentrations (shown in square brackets; μmoles/liter, i.e., μM). *V*_*c*_ and *V*_*e*_ are the intracellular volume of one cell and the volume of the culture medium accessible to one cell respectively. Bound and free transporters are expressed in terms of the amount present per cell (μmoles/cell). A mass balance on each species for transport involving *n*_*A*_ AAs, *n*_*T*_ transporters, and *n*_*A*_ × *n*_*T*_ possible AAT-AA complexes on each side leads to a coupled system of 2 × (*n*_*A*_ + *n*_*T*_ + *n*_*A*_ × *n*_*T*_) ODEs, summarized by equations 3-5.

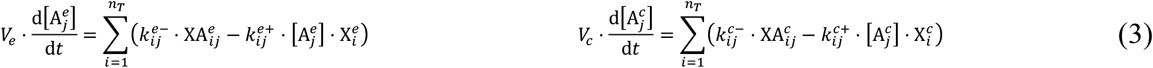

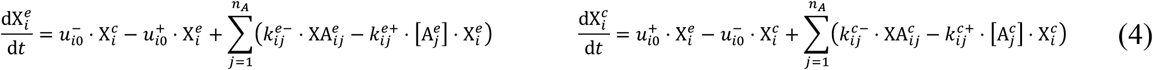

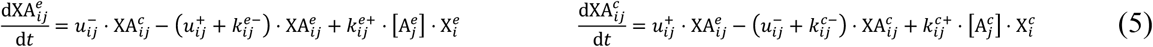

For each AAT-AA interaction, the full model requires six rate constants which can be approximated from *V*_*max*_ and *K*_*m*_ values as discussed in the methods section. Conservation of the free and bound transporters is included through an appropriate choice of the initial conditions. Equations (3) to (5), along with the initial conditions listed in SI, form an initial value problem (IVP) which can be numerically integrated. We have constructed a MATLAB tool that can generate the IVP automatically based on the details of AAs and AATs provided (see SI). Model parameter values (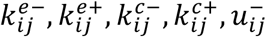 and 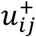) and the system ODEs are generated from the user-specified initial conditions, *V*_*max*_, and *K*_*m*_ values. The tool allows easy insertion or deletion of elements (AATs or AAs) of the transport network. In addition to these numerical solutions, equations (3) to (5) can be solved analytically for steady state for specific conditions. This model ignores the contribution and changes in the co-transported ions. The rate constants are not parameters for elementary reactions, and the steady state should not be interpreted as thermodynamic equilibrium.

### Simplified model for calculation of uptake rates

The uptake rates can be calculated by solving the system of equations for the time-varying values of the free and bound AA and transporter concentrations, but this approach is analytically intractable. The (pseudo)steady state uptake rates can be estimated by solving for concentration following the fast transients. Analytical expressions for the steady state concentrations can be derived (see SI) for several situations. For a 1-uniporter-*n*_*A*_AA system (*n*_*T*_ = 1, *n*_*A*_ > 1) at steady state, the ratio of external to internal AA concentrations is independent of the transporter level and affinity for uniporters. For a 1-exchanger-*n*_*A*_AA system, the steady state ratio of carriers on the two sides (*X*^*e*^: *X*^*c*^) is found to be proportional to the ratio of AAs on the two compartments 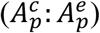 (see SI). It is however obvious that in practice, the steady state distribution will depend not only on AA transport but also on AA formation and utilization, as well as regulation of transporter level and activity.

The initial uptake rates are less dependent on metabolism and regulation. We have derived an analytical expression for the uptake rates for a simplified system with constant external and cytosolic AA concentrations. Such a system can be represented using four equations for each AA-transporter combination (details in SI). The rate of uptake of AA A_*p*_ can be quantified by the rate of translocation of carrier–A_*p*_ complex. The sum of net translocation rates of all complexes containing A_*p*_ gives the net uptake rate of A_*p*_.

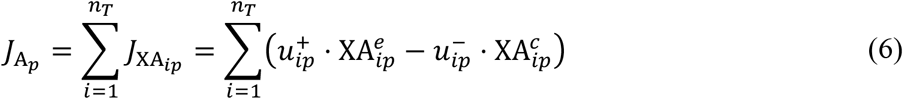

Algebraic expressions for free and bound carrier levels at steady state can be derived (detailed derivation in SI). Substituting the expression for 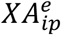 and 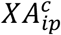 in equation (6) yields the net steady state uptake rates in terms of scaled AA concentrations.

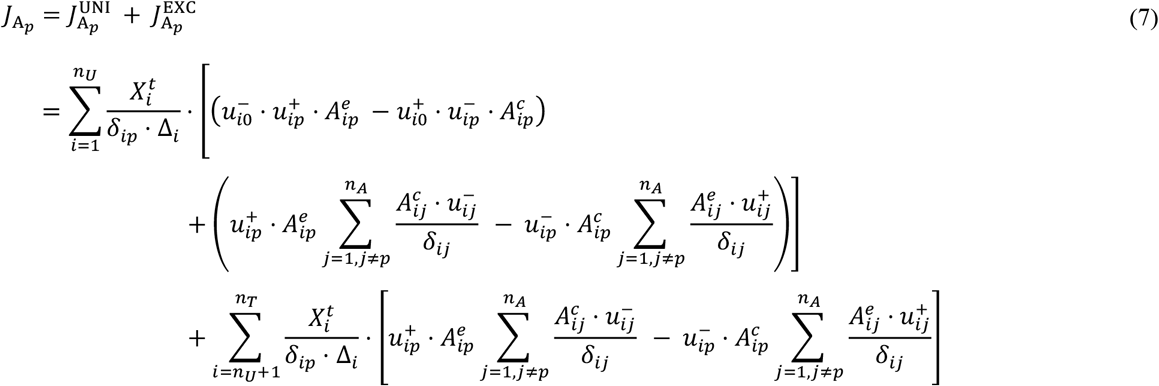

The terms 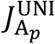 and 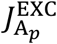 correspond to the net steady state influx of A_*p*_ through *n*_*IJ*_ uniporters and *n*_*T*_ − *n*_*U*_ exchangers. Here, 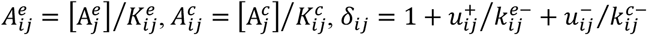, and

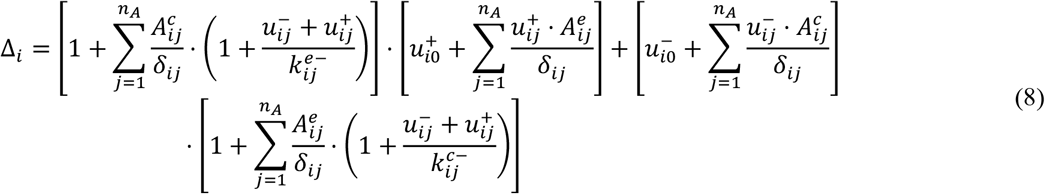

Since the AA concentrations are unvarying, the steady state solutions obtained here are equivalent to the pseudo steady state between AAs and carriers, as previously mentioned. Hence, the expression for 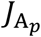 serves as an accurate descriptor of the initial uptake rates in the full model. If cellular concentrations for all AAs are assumed to be zero, the initial flux of amino acid A_*p*_ through a single transporter reduces to a form similar to the Michaelis-Menten equation for multiple alternate substrates.

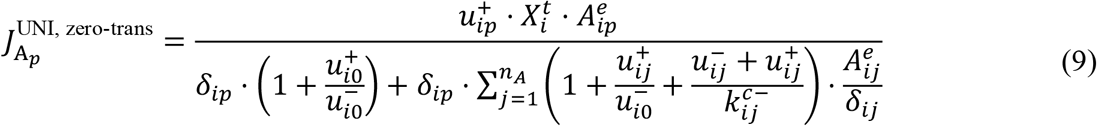

When the translocation rates are comparable, that is 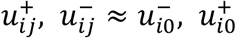, then the resulting equation is identical to competitive inhibition expression with *n*_*A*_ − 1 inhibitors. During a zero-trans experiment, the derived uptake rates are similar to the non-carrier-based model formulation, such as the Pardridge model. The simplified model was compared with the full model and existing models listed in Table 1 (details in SI).

### Development of a CHO-specific model

Based on reported AAT transcript abundances(Geoghegan et al, 2018), we have selected AATs with an expression level > 5 FKPM (Table 2). The 17 AATs thus selected were used to construct a detailed model of the CHO AA transport system (Figure 2, SI Figure 6). To our knowledge, this is the first comprehensive model for AA transport in CHO cells, or any mammalian cell type. *K*_*m*_ and *V*_*max*_ values are collected for each interaction shown in Figure 2, and are supplied to the MATLAB tool for calculating the model parameters and defining the system of ODEs as explained in the methods section. In addition to the kinetic parameters, transport specificity and direction for each AA has to be provided for calculating model parameter values (equations (1)-(2)). Initial conditions for transporters are determined based on the average expression levels shown in Table 1. All parameters used for simulation are listed in SI.

**Table 2:** List of AATs in the amino acid transport network in CHO cells. ‡ represents AAT + SLC3A2 complex.

| No. | Gene | Synonyms | Mechanism | Transport Specificity |  |  | Avg.<br>FPKM |
| --- | --- | --- | --- | --- | --- | --- | --- |
|  |  |  |  | Inwards | Outwards | Bidirectionally |  |
| 1 | SLC1a3 | EAAT1, GLAST | Na+ Sym. | Glu, Asp |  |  | 28.95 |
| 2 | SLC6a9 | GlyT1 | Na+ Sym. | Gly |  |  | 9.98 |
| 3 | SLC6a15 | B0AT2, SBAT1 | Na+ Sym. | Pro, Leu, Val, Ile, Met |  |  | 19.44 |
| 4 | SLC7a1 | CAT1, ERR | Uni. | Arg, Lys |  |  | 25.80 |
| 5 | SLC36a1 | PAT1, LYAAT1 | H+ Sym. | Pro, Ala, Gly |  |  | 9.98 |
| 6 | SLC36a4 | PAT4, LYAAT2 | Uni. | Trp, Pro |  |  | 14.10 |
| 7 | SLC38a2 | SNAT2, SAT2 | Na+ Sym. | Gly, Pro, Ala, Ser, Cys, Gln, Asn, His, Met |  |  | 84.65 |
| 8 | SLC38a6 | SNAT6, NAT1 | Na+ Sym. | Glu |  |  | 9.26 |
| 9 | SLC38a7 | SNAT7 | Na+ Sym. | Gln, Asn, His, Ala, Ser, Asp |  |  | 15.41 |
| 10 | SLC38a10 | PP1744 | Co-transporter |  | Ser | Gln, Glu, Ala, Asp | 117.12 |
| 11 | SLC43a2 | LAT4 | Uni. |  |  | Ile, Leu, Phe, Val, Met | 22.60 |
| 12 | SLC1a4 | ASCT1, SATT | Exchanger. | Ser, Thr | Asn | Ala, Cys | 89.74 |
| 13 | SLC1a5 | ASCT2, AAAT | Exchanger | Gln, Asn | Ala, Ser, Thr | Cys | 35.85 |
| 14 | SLC7a5<sup>‡</sup> | LAT1, 4F2lc | Exchanger | Leu, Ile, Val, Phe, Met, Arg, Trp, Tyr | Gln, Asn |  | 101.99 |
| 15 | SLC7a6<sup>‡</sup> | Y + LAT2 | Exchanger | Gln, Leu, Met | Arg, Lys, His |  | 17.71 |
| 16 | SLC7a7<sup>‡</sup> | Y + LAT1 | Exchanger | Gln, Leu, Met | Arg, Lys, His |  | 8.51 |
| 17 | SLC7a11<sup>‡</sup> | xCT | Exchanger | Cys | Glu |  | 13.26 |

**Figure 2:**
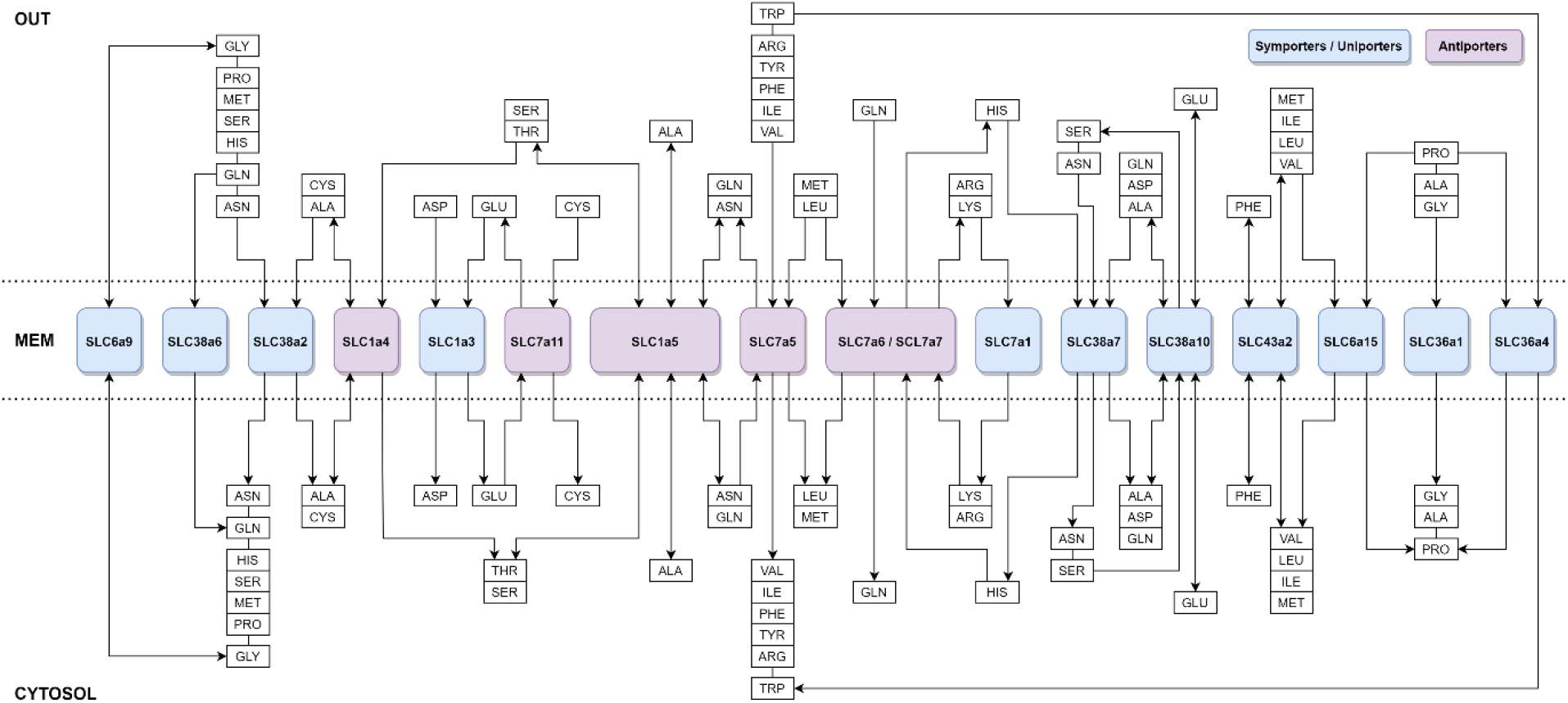
Amino acid transport system in CHO cells.

### Prediction of net flux from amino acid concentrations

In order to test the validity of the model, we compared model predictions of uptake flux to that reported in two different studies by different experimental groups, viz., Hansen-Emborg(Hansen & Emborg, 1994) and Schmid-Keller(Schmid & Keller, 1992). Neither of these studies were used in model construction or estimation of reaction rate and transporter abundance parameters. The initial uptake rates for each condition were calculated using two approaches as described in the Methods section, numerically for the full system and analytically for a simplified system assuming constant cytosolic and extracellular concentrations. It was found that the initial uptake rates obtained using equation (7) are identical to the results obtained from the numerical simulations. Hence, for further analysis, the initial uptake rates of AAs computed using the analytical method are presented.

Figure 3 shows the net initial uptake rates determined using equation (7) for the 16 AAs reported by Hansen-Emborg. Experimental data is for uptake by CHO cell lines in Phase II and Phase III after the initial phase one where the culture conditions changed from batch to continuous mode. It is seen that there is a qualitative match of the model predictions to the experimentally reported values for 12 out of the 16 AA fluxes, with several showing a quantitative match in both phases (Figure 3 top and bottom subplots). It should be noted that the transporter levels were estimated from an independent source, and the affinity and intra-membrane transport rate constants were also estimated independently. It is likely that the exact affinities and transporter levels may be different, corresponding to the culture conditions used by Hansen and Emborg. Four AA fluxes (alanine, arginine or asparagine, aspartate, and histidine) show qualitatively different uptake rates which may reflect an incorrect estimation of transporter levels or rate constants. Similar concordance between the model prediction and experimental uptake flux values were obtained for the Schmid and Keller data (see SI Figures 11-13) on AA concentrations and corresponding uptake flux values in murine hybridoma cells.

**Figure 3:**
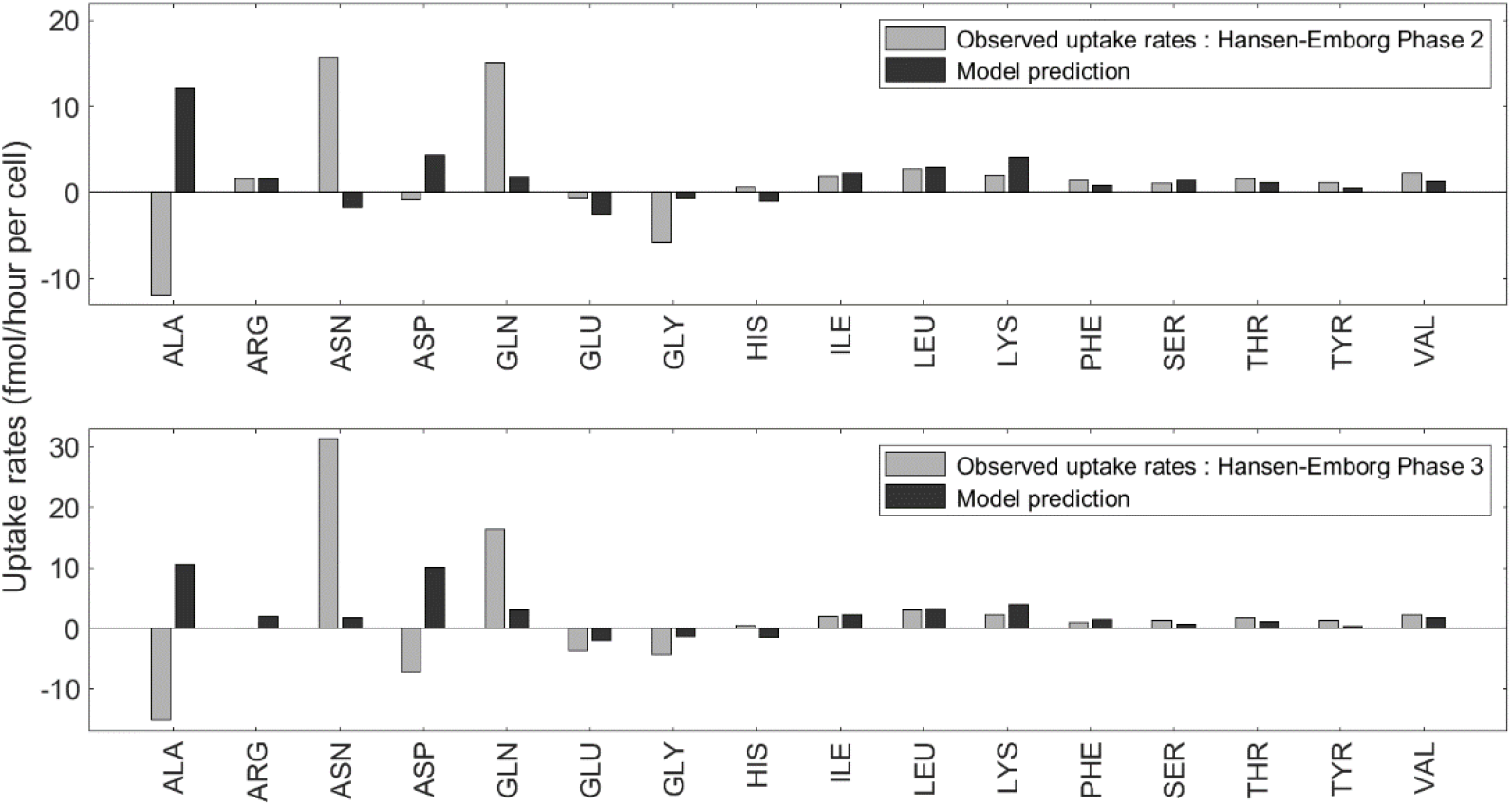
Amino acid initial uptake rates in CHO cells calculated analytically for the simplified model based on equation (7) for cellular and external AA concentrations corresponding to Phase II (top) and Phase III (bottom) Hansen-Emborg data.

### Prediction of change in flux due to amino acid and transporter perturbations

Several reports on mammalian cells (not limited to CHO) have listed the effect of specific AA perturbations on the uptake rate of other AAs. We have used the analytical expression for uptake rates (equation (7)) to qualitatively estimate the effect of perturbations in AA and transporter levels by calculating the partial derivatives of uptake rates (equation (7)) with respect to external and cellular AA concentrations.

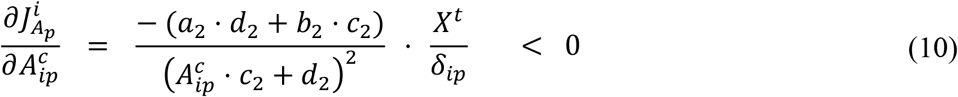

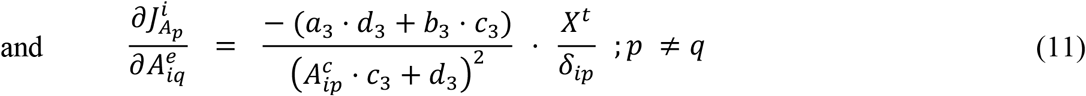

Here *a, b, c, d* (see detailed equations in SI) are combinations of AA concentrations and kinetic parameters that are always non-negative. From equation (10) it is clear that the cellular addition of an AA always decreases its initial uptake rate, and external addition of AAs increases the uptake (See SI). This result is independent of the transporter types and concentrations of other AAs. In contrast, the rate of change of uptake rate of *A*_*p*_ on the addition of another amino acid 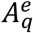 (equation (11)) depends on the term *b*_3_, which can be either positive or negative depending on the values of concentrations and kinetic parameters. If *b*_3_ > 0, addition of 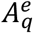 will decrease the uptake of *A*_*p*_. Results of perturbation analysis calculating the effect of a change in external AA concentration on all AA uptake rates for the Phase II Hansen-Emborg intracellular and extracellular AA concentrations are shown in Figure 4. For the cells along the principal diagonal, it is seen that the external addition of an AA resulted in flux changes greater than one for the AAs that have an existing positive flux (lightly shaded cells in figure 4) and a value less than one (decreased efflux) including negative values (reversal of the efflux to influx) for cells that have existing efflux (dark cells). In brief, the increase in external AA concentration results in an enhancement in its initial uptake rates (or decrease in efflux rates) for all AAs, which agrees with the partial derivatives derived earlier. Conversely, cellular addition mitigates the intake.

**Figure 4:**
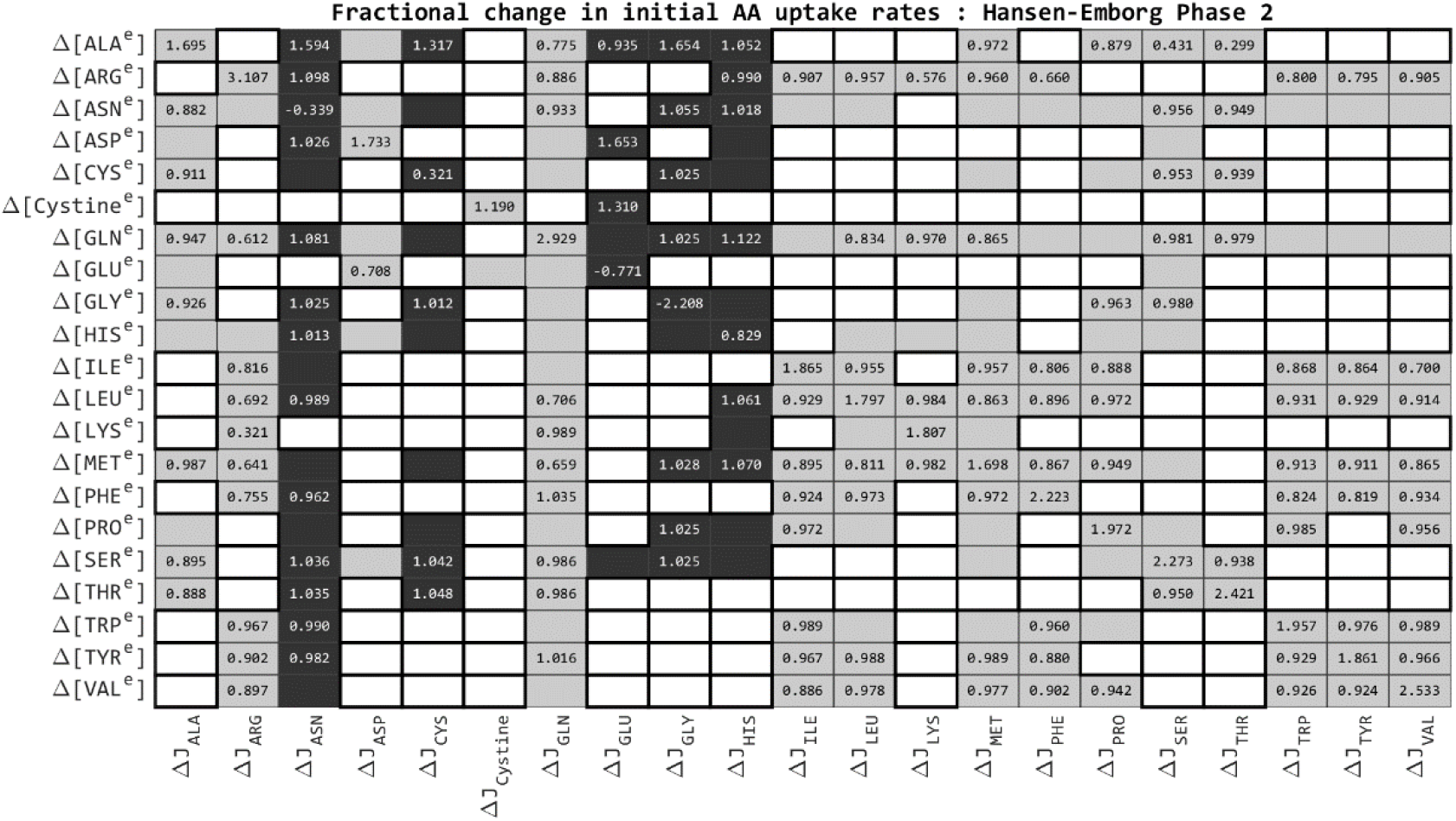
Effect of external AA concentration changes on initial uptake rates. Each external AA concentration was increased by 100%, and the resulting fold-changes are calculated. AA pairs that do not share a common transporter are highlighted in white squares. Empty shaded cells represent values below the threshold, 0.01. Light grey and dark grey cells indicate initial transport direction, into and out of the cell respectively.

Significant off-diagonal values represent connected AA pairs that share multiple transporters. Glutamine-asparagine is one such interdependent pair. Both glutamine and asparagine are used as exchange substrates for many other AAs in exchangers such as SLC7a5 and SLC1a5, and the supplementation of either of these AAs inhibit the uptake of the other.

### Model predictions are consistent with reported amino acid interdependencies

Specific predictions of the effect of AA perturbations on flux are consistent with experimental reports in other cell types. For instance, in liposarcoma and breast cancer cells, it has been shown that Asn preloading in LPS2 cells led to significantly higher intracellular Ser and Thr levels, and a modest increase in Leu/Ile, Met, Phe, Trp, Arg, Lys and Asp. There was no effect on Gly and Pro(Krall et al, 2016). Using equation 7, the effect of a 100% increase in Asn from the Hansen-Emborg Phase II levels on the uptake flux of all AAs was calculated. The simulations predict enhanced Gln, Ser, Thr, Arg, Phe, Trp, Ile, Leu and Met uptake rates. The same trend is also observed if Phase III data is used, though the effect is lower. Effect on Lys uptake is zero since this CHO-specific network does not contain any common transporter for Lys and Asn. The model predicts that Gly/Pro uptake is not affected, consistent with the experimental report.

Simulations were carried out corresponding to experimental reports of inhibition of threonine uptake by Cys, Ser, Ala, Leu in a kidney epithelial cell line BSC-1(Kuhlmann & Vadgama, 1991). Model predictions also predict inhibition of threonine uptake following external addition of Ala, Cys and Ser. In addition, the model also predicts an inhibition following Asn addition, but fails to predict the observed response to Leu since there is no common transporter in the CHO network used for the simulations.

Similarly, model simulations corresponding to experimental reports of inhibition of valine uptake by leucine(Torres et al, 1995), glutamate uptake by aspartate(Drejer et al, 1982), and proline uptake by glycine, serine and alanine(Nicklin et al, 1992) were carried out using Phase II/III AA concentrations and the CHO AA transport network, and found to be consistent with the experimental observations. This ability to predict AA interactions across (mammalian) cell lines suggests that the AA transport network is largely functionally similar across different cell types. However, a definitive analysis will require construction of cell-type specific networks based on transporter expression data for each cell line.

## Discussion

The generalized model presented here is to our knowledge the only model that can simulate the kinetics of individual and net AA uptake by an interconnected system of multiple AAs and transporter types, while explicitly accounting for the differences in AAT-AA interactions. Identical rate constants for all AAT-AA interactions are appropriate while considering isotopic forms or labeled forms of the AAs, but several studies indicate that each AA interacts differently with AATs. Hence rate constants for AA-AAT interaction should be specific to each AA-AAT pair. This model includes the contribution of obligatory exchangers and uniporters to the overall transport rate. Through a suitable choice of binding and transport parameters in the uniporter model, or by modelling the concurrently transported ion’s counterion in an exchanger model, it can also be used to model symporters. The MATLAB tool automates the process of creating IVPs for any specified transporter profile and AA concentrations. The MATLAB tool can be easily customized to simulate different media, cell types and initial conditions.

From the initial concentrations, kinetic parameter values (*K*_*m*_, *V*_*max*_), and information on the direction of AA transport through each carrier (*K1* and *K2*), the full model is able give a comprehensive picture of the AA transport process under different extracellular and intracellular conditions. We have used expression data to estimate transporter levels. However for some transporters, transcript levels do not correlate to protein levels(Gauthier-Coles et al, 2021).

Information on direction of transport and transporter type is not always unequivocal. For instance, SLC38a10 is classified as a bidirectional transporter and an ion exchanger(Geoghegan et al, 2018; Hellsten et al, 2017). Values of specific reaction rate parameters are not always available and approximations have been used to estimate these values as described in the methods section. Therefore, the model predictions have to be regarded as estimates of the quantitative uptake rates. Nevertheless, it is seen from Figure 3 (and SI Figures 11-13) that there is a quantitative match with the measured experimental values for many AA uptake rates. Given that the experimental data was not used for defining the transport network or its parameters, this concordance across cell lines and cellular/external AA compositions serves as a validation of the model’s utility.

The steady state concentrations calculated using this model are clearly of relevance only in the context of a reconstituted system or an in vitro assay. AA metabolism is not part of this model, so the calculated steady state concentrations and rates do not account for AA interconversion, formation, or incorporation in the cell’s central metabolism. The interphase transfer of AAs across the distinct extracellular and intracellular environments is approximated by a carrier model. The individual reactions are not elementary since they omit the contribution of ions and dependence on factors such as pH. As such the calculated and analytical stationary states are steady state and not equilibrium concentrations. Besides, it is known that the transporter levels are regulated(McGivan & Pastor-Anglada, 1994), and therefore unlikely to remain constant over long time scales.

Initial uptake rates computed for the simplified system (equation (7)) help dissect the effect of substrate competition in the AA transfer. The expression for the net uptake through uniporters 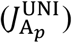 displays a clear separation between self and interaction effects of amino acid A_*p*_. The self-effect 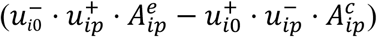 depends on the rate of free carrier translocations. This rate can also be used as a qualitative measure of the effect of substrate competition on the uptake of AAs. With a high translocation rate of uniporter, the self-effect term dominates the expression for net initial uptake. As 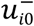 and 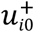 decrease, the impact of substrate competition on the uptake rates increases. Net initial uptake through exchangers, on the other hand, contains only the interaction term since the exchange process requires the interaction of multiple AAs with the exchanger. Consequently, in a mixed system of uniporters and exchangers, very high translocation rates of unbound uniporters can lead to the repression of transfer through exchangers.

Numerical simulations of the CHO cell AA transport network show the effectiveness of the model (and the MATLAB tool) in studying the effects of different culture media conditions and cellular AA profiles on the uptake of AAs. Although the perturbation analysis is carried out using two methods, numerically by comparing the initial uptakes before and after perturbation and analytically by computing the partial derivatives of uptake rates, the qualitative nature of the results obtained are identical, attesting to the utility of the simple calculation which avoids the process of numerical integration of the full model.

Partial derivatives of the uptake rates with respect to external AA concentrations (equations (10) and (11)) suggest that AAs that are not directly connected do not influence each other’s initial uptake rates. Such indirect dependencies may manifest at a later time. Using numerical simulations, it was verified that the non-effect between unconnected AAs persists for a long (∼1 hour) time period and suggests that this analysis may be relevant for the initial uptake rate, not just the instantaneous rate (details in SI). In the case of AAs that share at least one common carrier, the full model predicts that the effect of perturbation could be observed at a later time point.

The model predicts that the external addition of an AA always results in the enhancement of its uptake rate (equation (10)). Based on substrate competition it is expected that the addition of an AA inhibits the uptake of its competitor and can be verified via numerical simulations. However, the partial derivative of uptake rates with respect to the concentration of competing AAs indicates that an increase in the uptake rate of one AA upon increase in the external concentration of another could also be observed for a suitable set of initial conditions and kinetic parameters (see SI for discussion of the role of parameter b3 in defining the sign of the numerator of equation (11)). The heat map (Figure 4) helps identify associations between AA pairs in the transport network. Simulation of the effect of instantaneous knockout of transporters on AA flux (Figure 18 in SI) shows the utility of the model in identifying significant (and insignificant) transporters in the network. An absence of significant effect on the transport of the AA being transported by that particular transporter indicates redundancy in transport. For instance, SLC36a4 knockout affects only a single AA, in contrast to the other AATs, whose influence is seen on almost all AA transport rates. Although SLC6a9 (GlyT1) transports glycine in both directions, the model predicts that the removal of GlyT1 will result in the enhancement of net glycine uptake.

The proposed model serves as a tool to identify and quantify the interrelationships between AAs arising from the topology of the transport network. The predictions of the AA uptake rates will also be useful to estimate AA requirements not only for other cell types but even for cell free synthesis systems(Bundy et al, 2018). The simplified model also enables the numerical and analytical study of the relationship between the culture medium conditions and AA transport. By determining the rate of transport of each AA through each carrier, significant amino acids and carriers in a transport network can be easily identified and their influence on other components can also be evaluated. The model can serve as a framework for interpreting uptake data for the same cell type under different extracellular conditions, or for different cell types. In addition, this model can be coupled to genome-scale metabolism/expression models(Salvy & Hatzimanikatis, 2020) or amino acid metabolism models(Kontoravdi et al, 2007) and serve as the transporter dynamics component of a comprehensive model for simulation of intra- and extra-cellular amino acid dynamics in different cell lines.

## Methods

### Model formulation and simulation

A general model of AA transport (henceforth referred to as the full model) was constructed based on the reaction network depicted in Figure 1. Equating the change in concentration to the net rate of association, dissociation, and transport resulted in the formulation of a system of ordinary differential equations (ODEs) representing the mass balance for each species involved in the transport process (equations (3)-(5), also see SI). MATLAB (version R2017a) function *ode15s* was used for numerical integration of the system of ODEs. The results are then compared with different limiting cases, as well as for exchanger and uniporter facilitated transports, both analytically and through numerical simulations. For constant cellular and external AA concentrations, an analytical solution for the steady state AA transport rate for a general *n*_*T*_AAT-*n*_*A*_AA system is derived, and used for computing the initial AA uptake rates. The full model was compared with the existing models of membrane transport using percentage deviation at individual times and RMS distance between the predicted time series (details in SI).

### Model and parameters for amino acid transport in CHO cells

The AA transport network for CHO cells was constructed based on transcriptome data for AA transporters(Geoghegan et al, 2018). We have assumed that mRNA expression levels correlate with protein levels, and that AATs with lower expression levels do not significantly contribute to the overall AA transport. 17 AATs with expression levels (FPKM) greater than 5 were used to construct the AAT network. Three AATs (SLC3a2, SLC6a6, and SLC43a3) are not included as they are reported to only act as enhancers for other AATs or do not carry a substrate of interest (Furukawa et al, 2015; Jhiang et al, 1993; Torrents et al, 1998). The transport of 20 AAs through 17 AATs in CHO cells, when represented as carrier-mediated transport processes, gives rise to 222 interacting species. A MATLAB script was written for generating the mass action rate laws and systems of ODEs based on the interaction network provided. The effect of the initial transient due to AA-AAT binding was eliminated by allowing the transporters to reach pseudo-steady state. Pseudo steady state concentrations and uptake rates were determined both through numerical integration and using the analytical expression derived for the simplified model. Positive uptake rates indicate a net transport into the cell.

CHO cell volume was approximated as 1.63 × 10^−12^ mL, consistent with the reported cell diameter range of 14.02 to 15.21 μm(Han et al, 2006). Assuming a cell density of 10^6^ cells/mL, the external volume accessible to one cell is estimated as 10^−6^ mL assuming that the volume occupied by cells is small compared to the total volume. The Michaelis-Menten constant (*K*_*m*_) and maximum transport velocity (*V*_*max*_) values have been measured for several individual AAT-AA systems. The available values for the 17 selected AATs were collected from different studies, and the remaining were approximated from the known values for similar AAs/ transporter types. If a similar AA or AAT is unavailable, the required values were taken as the average over all uniporters or exchangers. The details of the parameter values are given in SI.

The model kinetic parameters (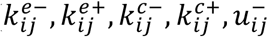 and 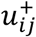 were approximated from the *K*_*m*_ and *V*_*max*_ values. *V*_*max*_ is assumed to be proportional to the translocation rate of the carrier–substrate complex. Superscript or subscript *ij* indicate that the constant is associated with the transport of AA A_*j*_ by the carrier X_*i*_. We define

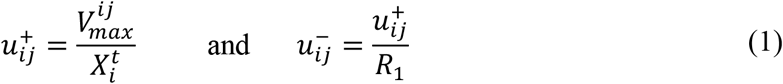

where 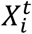 and 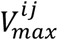 respectively denote the total amount of the carrier X_*i*_ present per cell and the known *V*_*max*_ for the transport. If the AA is instead known to be released by the cell, then the value of 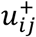 will be interchanged with 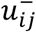. The dissociation rate constants 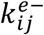 and 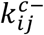 for all AAT_*i*_ − AA_*j*_ pairs were set to 1000 min^−1^ based on dissociation values for receptors and symporters(Lauffenburger & Linderman, 1996; Mim et al, 2005). The association rate constants were then calculated such that if the transport is inwards,

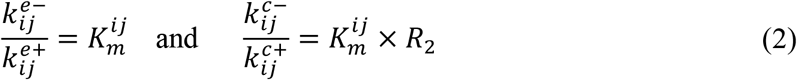

If the transport direction is outwards, then the value of 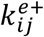 is exchanged with 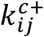. As no information is available on the translocation rates of the free carrier, both 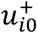 and 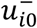 were fixed to 10^6^ min^−1^, consistent with the assumption that the unloaded carrier translocation is not rate-limiting. The terms *R*_1_ and *R*_2_ determine the extent of directionality imposed on transport. As such, the model can represent both uniporters and symporters without needing the inclusion of the co-transported ion that drives the unfavorable flux in the case of symporters. We have assumed *R*_1_ = 10 and *R*_2_ = 100. In the case of the AAs that are transported bidirectionally (Table 2), the parameters are assumed to be symmetric (i.e., *R*_1_ = *R*_2_ = 1).

Analysis of CHO RNA-Seq data(Singh et al, 2018) revealed that the highly expressed AATs, chosen as the CHO cell transport network representatives, correspond to 20% of the total membrane transporters. In a cell, there are approximately 6.64 × 10^−17^ moles of membrane transporters (Itzhak et al, 2016). The amount of each carrier on the membrane was calculated based on their relative expression levels. Initially, all AATs were assumed to be equally distributed on the cytoplasmic and extracellular sides. Based on numerical simulations, the time to reach a pseudo-steady state on the membrane surfaces is determined, and the corresponding AAT distribution is regarded as the initial condition for the AATs on the two surfaces. This pseudo-steady state is utilized as the initial condition for the numerical simulations.

To test the model, we have compared the model prediction of uptake rate with experimentally measured uptake rate and AA concentration data from two studies from different groups(Hansen & Emborg, 1994; Schmid & Keller, 1992), neither of which was used in model construction or calibration. For each simulation, extracellular and intracellular AA concentrations were set at the experimentally reported values, and the uptake flux was calculated using the simplified model. Transporter initial conditions are assumed to be the same for all AA initial conditions (see SI for details). The model simulation/calculation included all 20 AAs, but only the 16 AA uptake rates that are reported by Hansen-Emborg are shown in the results.

The effects of the CHO cell AA transport network on the uptake and efflux rates of AAs are evaluated by perturbing the extracellular AA concentrations and transporter expression levels and recording the changes in initial uptake rates. Perturbation analysis was performed by numerically integrating the model and evaluating the changes in the initial uptake rates or concentrations of AAs at 1, 10, and 100 mins. In all cases, the initial conditions were set to the pseudo steady state values following AA-AAT binding. Initial uptake rates of AAs following perturbation to AA/carrier levels are also computed analytically for the simplified system with constant AA external and cytosolic levels. The partial derivatives of uptake rates were derived and used to quantify the effect of each species on the initial uptake rates.

## Data availability

The information (equations, parameter values) used to obtain the figures are completely specified in the Methods section and Supplementary Information file. Computer code used for generating the figures is available from the corresponding authors upon reasonable request.

## Acknowledgements

This work was supported through grants from the Science and Engineering Research Board, India to MG (CRG/2020/003869) and CG (EMR/2017/003271). We thank Harsh Tiwari, a NCL summer intern from the School of Biochemical Engineering, Indian Institute of Technology (BHU) for help with the literature survey for model parameters.

## Author contributions

Conceptualization: M.G.; Model formulation: A.S., M.G., C.G; Mathematical analysis: A.S., C.G; code and simulations: A.S.; analysis of results: A.S., M.G., C.G; resources, supervision: M.G., C.G.; writing: A.S., M.G., C.G

